# COLT-Viz: Interactive Visualization of Antibody Lineage Trees

**DOI:** 10.1101/219592

**Authors:** Chenfeng He, Ben S. Wendel, Jun Xiao, Keke Chen, Ning Jiang

## Abstract

Many tools have been developed to visualize phylogenetic trees, which is a traditional technique for evolutionary tree analysis. However, due to the unique characteristics of antibody lineage trees, the phylogenetic method cannot adequately construct proper tree structures for antibody lineages, and many other tools have been developed to address this problem. However, there still lacks of an adequate tool to visualize the resulted antibody lineage structures that are more complicated than phylogenetic trees. In addition, high-throughput sequencing-based antibody repertoire profiling enables the counting of the number of transcripts associated with individual antibody sequences, thus more dimensions need to be encoded in the tree structure visualization. Further, users may wish to manually adjust the tree structure for a special context. When doing so, they may wish to maintain some biological constraints that are applicable in antibody lineage tree structure, such as isotype switching constraints or different sampling constraints. Here, we report an interactive visualization tool (COLT-Viz) designed to display the number of RNA copies, number of somatic hypermutations, and sample collection time associated with each antibody sequence as well as the distance to neighboring sequences for each antibody sequence in the lineage. COLT-Viz also allows users to interactively visualize and edit antibody lineage structures while giving users the option to automatically check biological constraints on the edited structures to ensure accuracy. COLT-Viz takes JSON text format as input files and can easily be used to visualize networks with or without the biological constraints. We believe the amount of information that can be displayed for complex antibody lineages, the interactive interface, and the option of checking for biological constraints make COLT-Viz a versatile tool for antibody lineage tree visualization that will guide further biological discoveries.

## 1. Introduction

Lineage analysis is frequently used by researchers to analyze the evolution of different species of organisms or to trace cellular development within an organism. Specifically in the immune system, lineage analysis has been adopted to trace the mutational evolution of antibodies (1–3) and understand selection pressure (4,5).

Typically, an antibody lineage is represented as a tree structure, which describes the possible development path (i.e., edges in a tree) among all unique antibody sequences (i.e., nodes in a tree). Traditionally, researchers used phylogenetic trees for antibody lineages; however, a phylogenetic tree is a binary tree with all nodes presented as leaf nodes, which is not suitable for antibody lineages (3). Much work have been done to construct tree structures specific to antibody lineages (3,6,7); however, there is still a need for a tool to efficiently visualize these trees. The visualization of antibody lineage trees has a few unique needs: (a) there are multiple features specific to antibody sequences (e.g., number of transcripts, number of somatic hypermutations, isotype, etc), integrating all these properties together for tree visualization will facilitate lineage analysis; (b) interactively visualize the antibody structure by showing antibody sequence information and associated properties when users click on a node of interest. This allows user to have access to detailed information of nodes (antibody sequences) without losing the global view of a complex antibody lineage structure; and (c) interactively modify antibody lineage structure. Oftentimes, multiple valid lineage tree structures exist for an antibody lineage. Based on special considerations, researchers may want to manually alter their tree structures by changing or adding connections between nodes. When doing so, they may want to make sure that these modified connections do not violate certain biological constraints. These constraints can either be biological constraints (8) or constraints determined by experiment design (9).

Although many network visualization tools with powerful functions have been developed (e.g., Cytoscape (10), Graphviz (11)), they were not designed for antibody lineage structure visualization and thus lack the consideration of biological properties and constraints. To our knowledge, there is no tool specifically designed to visualize antibody lineage tree structures with the functionalities mentioned above. For example, LymAnalyzer (12) incorporates FigTree (13) to visualize antibody lineage tree structures. However, FigTree is only designed for phylogenetic tree which is not suitable for antibody lineage structure and does not have an interactive user interface. Another program, Change-O (6), modifies phylogenetic tree analysis for antibody lineage application, but its visualization function can only generate non-interactive figures. In addition, none of these tools can easily be used to visualize the multiple features of antibody lineages, which are unique to antibody lineage tree visualization.

Bearing the drawbacks of current visualization tools in mind, we developed COLT-Viz for antibody lineage tree structure visualization. COLT-Viz enables researchers to interactively explore lineage trees with multiple antibody sequence properties encoded and automatically checked when the tree structure is manually altered. At the same time, researchers have the option to override error messages and proceed with the desired changes. This tool enables users to discover interesting patterns of antibody evolution process and select interesting nodes (antibody sequences) for further evaluation.

Specifically, COLT-Viz projects antibody sampling time, total mutations for a sequence, mutation distances between two neighboring sequences, and antibody RNA copies of unique sequences onto different elements of each node in the lineage visualization, which allows users to intuitively understand the lineage evolution over time, mutation distances from a germline sequence, mutation distances between neighboring nodes, and the clonal size associated with each unique antibody sequence. Furthermore, users can fine tune the lineage by removing edges and reconnecting nodes with all the constraints automatically checked by the system with an option to either override the error message or make other alternative changes. When users edit the connections, COLT-Viz checks two types of constraints: (a) isotype constraint: as described in previous studies (8,14–16), antibody isotype switch can only proceed according to the order of IGH constant genes located on the chromosome (17), thus users may want to maintain this constraint while editing the lineage; (b) time constraint: as described in Schramm et al. (9), longitudinal information offers another dimension in some biological experiments (18). Although in many cases, sequences from an earlier time point may not necessarily be ancestors of sequences from later time point, in some cases, such as transplantation, where donor samples and recipient samples are experimentally separated, this constraint is critical. Thus, users may want to re-enforce the time constraint when manually changing node connections. Therefore, COLT-Viz also checks if any manual changes on the tree structure comply with the time constraint that sequences from an earlier time point should be ancestors of sequences from a later time point while give users the option of override this error message and proceed with their intended changes.

## 2. Implementation

COLT-Viz takes the lineage tree from tree-generating tools (e.g., COLT (7), Change-O (6)) as input and outputs the tree visualization. The input of COLT-Viz contains one node file and one edge file. The node and edge files are encoded in JSON (JavaScript Object Notation) format (http://json.org/), which is a lightweight and human-readable text format. JSON expresses data objects in attribute-value pairs, which makes it an ideal data format for visualizing antibody sequences with multiple attributes. Outputs from tree-generating tools/algorithms can be converted to the JSON text format to be compatible with COLT-Viz. Users can also manually edit the JSON text files to change the lineage tree structure. The node file contains a list of nodes, or unique antibody sequences. Each node element includes the node identity, node description, associated sequence abundance (NRNAs, number of RNA copies, and NREADs, number of sequencing reads), antibody isotype (i.e., antibody heavy chain families, IgM, IgD, IgG, IgA, and IgE), timestamp (i.e., sampling time point, for example, when doing research on vaccine, sequences obtained before vaccination are all assigned to a smaller number (e.g. 20) while sequences obtained 7 days post vaccination are all assigned to a larger number(e.g. 40)), and the number of somatic hypermutations from the germline sequence. Each edge in the edge file is defined as a 3-tuple (parent node identity, weight, child node identity), which are also encoded in the JSON format.

Users can explore the tree visualization, e.g., checking the detailed information of each node, and fine tune the tree structure by editing and moving the edges. Before making changes, COLT-Viz automatically checks the constraints and displays a warning message if either the timepoint or isotype constraint is violated. However, user can choose to override this warning message and proceed with the move. Once old edges are removed and new edges are added, the system will automatically update the entire graph. Finally, the updated tree structure can be saved back to the edge files in the JSON format for later usage.

## 3. Results

### 3.1 Lineage Tree Visualization

We visually encode multiple properties of nodes and edges into the lineage tree visualization, so users can understand these properties intuitively. Figure 1 shows an antibody lineage we found in a young African child who experienced acute malaria twice in two consecutive malaria seasons from our previous study (19). There are a total of 23 unique sequences obtained from 4 timepoints in this lineage. The height of each node represents the number of RNA copies associated with that unique sequence. Heatmap is used to color nodes – the warmer the color, the greater the number of somatic hypermutations to the germline sequence. The border of each node is colored to represent different timestamps, so users can easily identify the evolution of the antibody lineage over time.

**Figure 1.**
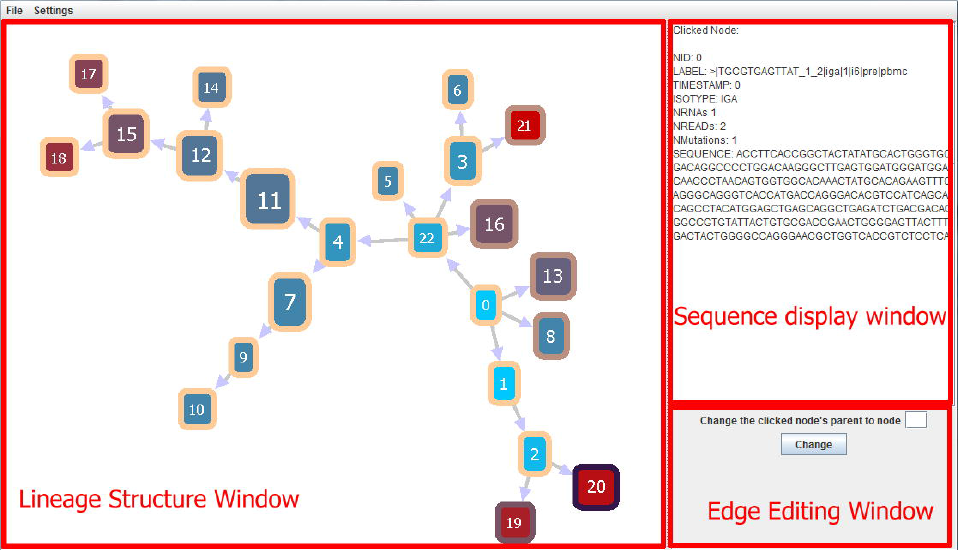
User interface of COLT-Viz. Within the ‘Lineage structure window’, height of each node represents the RNA copy of each sequence, color of each node represents the mutation with respect to the germline, color of the border of each node represents the time point and the edge length represents the edit distance between neighboring nodes if user specifies the option. By clicking on a node, the information of this node will be displayed in the ‘sequence display window’. User can also edit the connections through ‘edge editing window’. The ‘lineage structure window’ shows an antibody lineage found in a young child who experienced two consecutive malaria infection. 23 sequences were found from 4 time points and visualized using COLT-viz. Four time points are: year 1 pre-malaria infection (light orange node frame for most of the nodes displayed), year 1 acute malaria infection (brown node frame for node 16, 13, and 8), year 2 pre-malaria infection (light purple), and year 2 acute malaria infection (dark purple). Sequence 0 is the root and arrows point to the progeny sequences. Sequences from later time exist in the outer leaves of the lineage, which represents their evolution overtime.

The tree layout algorithm uses the well-known force-directed graph drawing algorithm (20). It progressively changes the nodes’ positions, as if there are physical forces between the nodes. Imagine that neighboring nodes are connected by springs: the force of the spring will prevent the nodes from drifting too far apart or collapsing too close together. This algorithm ideally places the root antibody sequence towards the center of the visualization and unfolds the tree outward to maximize the use of the display area. Thus the directionality of the tree is in general from center to periphery. Knowing this makes it easier for user to identify the root node, also if user assign the root as a specific ID (e.g., 0, same as in Figure 1), then the root node should be easily identified in the tree. Then, by following the directionality, users can easily understand the development of the whole lineage.

Using COLT-Viz, features of the antibody lineages that are not obvious by mining the data can be quickly perceived visually. Using the lineage in Figure 1 as an example, users can quickly identify the root sequence and see that, for this lineage, the sequences at later time points have accumulated more somatic hypermutations compared to sequences collected at earlier time points, and they lie in the outer leaves of the lineage. In addition, the relatively even distribution of node heights indicates that the RNA copies of these unique sequences do not differ widely. These visual features help immunologists quickly glean an overview of the evolution of this antibody lineage over time.

### 3.2 Tree Editing and Constraint Maintaining

COLT-Viz allows users to alter the network connections by removing and adding edges between nodes. When a new edge is added, the system will check two types of constraints: (a) time constraint, an edge x->y is valid only if x’s timestamp is earlier (smaller) than y’s; and (b) isotype constraint, an edge x->y is valid only if the isotype is either unchanged or following the class-switching rule: a class-switched sequence cannot switch back to IgM. If either of the constraints is violated, COLT-Viz will display a warning. However, users can override this warning and proceed to add the new edge or heed the warning and cancel the new edge. The constraint checking helps avoid artificial mistakes and maintain interesting time features when users manually modify antibody lineage trees.

### 3.3 Intuitive graphic visualization user interface

COLT-Viz employs an intuitive graphic visualization user interface (Figure 1). Within the antibody lineage structure displaying window, users can drag on any node to reposition the tree. Users can also click on any node to view the full antibody sequence and annotated information in the sequence display window on the right. In addition, users can change the lineage structure by clicking on a node and specifying the new parent node that they wish to establish a link in the edge editing window.

## 4. Discussion

COLT-Viz is an interactive tool designed for antibody lineage tree visualization and editing. By using COLT-Viz, researchers can understand lineage trees, validate algorithmic results, and fine-tune the tree structures with additional domain knowledge that automated algorithms cannot capture. Further, COLT-Viz allows researcher to identify nodes (antibody sequences) that bear specific features that are only obvious in graphic settings. Constraints are automatically checked to avoid manual editing errors. Although COLT-Viz is specifically designed for antibody lineage analysis, we believe with small tweaks it can be easily applied to any other type of lineage analysis in biomedical research.

## Availability of data and software

Project name: COLT-Viz

Project home page: https://github.com/immudx/paper (sample data is included).

Operation systems: Platform independent

Programming language: Java

Any restrictions to use by non-academics: None

## Abbreviations

IgM (D,G,A,E): Immunoglobulin M (D,G,A,E)

## Author contributions

NJ conceived the idea and directed study; CH, BW and NJ participated in the algorithm design. KC and JX wrote the computer software. CH, BW contributed to software testing and optimization. CH, KC, JX and NJ contributed to the writing of the manuscript. All of the authors read and approved the final manuscript.

## Acknowledgements

Not applicable

